# Genome assembly of the maize inbred line A188 provides a new reference genome for functional genomics

**DOI:** 10.1101/2021.03.15.435372

**Authors:** Fei Ge, Jingtao Qu, Peng Liu, Lang Pan, Chaoying Zou, Guangsheng Yuan, Cong Yang, Shibin Gao, Guangtang Pan, Jianwei Huang, Langlang Ma, Yaou Shen

## Abstract

Heretofore, little is known about the mechanism underlying the genotype-dependence of embryonic callus (EC) induction, which has severely inhibited the development of maize genetic engineering. Here, we report the genome sequence and annotation of a maize inbred line with high EC induction ratio, A188, which is assembled from single-molecule sequencing and optical genome mapping. We assembled a 2,210 Mb genome with a scaffold N50 size of 11.61 million bases (Mb), compared to those of 9.73 Mb for B73 and 10.2 Mb for Mo17. Comparative analysis revealed that ∼30% of the predicted A188 genes had large structural variations to B73, Mo17 and W22 genomes, which caused considerable protein divergence and might lead to phenotypic variations between the four inbred lines. Combining our new A188 genome, previously reported QTLs and RNA sequencing data, we reveal 8 large structural variation genes and 4 differentially expressed genes playing potential roles in EC induction.

**Highlight:** Our manuscript presents a high-quality reference genome of the inbred line A188, and provides new insights into candidate genes underlying maize embryonic callus induction and other maize agronomic traits.

## Introduction

Genetic transformation has been an effective approach for elucidating gene functions in plants. Maize (*Zea mays* L.) genetic transformation is highly relied on the utilization of embryonic callus (EC) induced from immature embryos. However, only a few lines possess the ability to efficiently form embryonic callus, including several inbred lines as A188, B104, H99, C01 and the combination Hi-II (A×B) etc. (Armstrong *et al*., 1992; Bronsema *et al*., 1997; Krakowsky *et al*., 2006; Salvo *et al*., 2018). Since the plant regeneration from maize tissue culture was firstly reported in 1975 (Green and Phillips, 1975), little is known about how the maize EC was induced from the immature embryos even though great efforts have been made by generation to generation of researchers (Armstrong *et al*., 1992; Krakowsky *et al*., 2006; Pan *et al*., 2006; Salvo *et al*., 2018). To date, only several QTLs were identified involved in controlling callus induction or plant regeneration.

Using an F_2_ population derived from two inbred lines with divergent EC induction rate, Pan et al. mapped 5 QTLs for tissue culture response on chromosome 1, 3, 7 and 8, respectively (Pan *et al*., 2006). Similarly, several type I callus formation related QTLs were identified using an F_6_ RIL population constructed from H99 and Mo17 (Krakowsky *et al*., 2006). More QTLs responsible for plantlets and transformation were identified using a segregation population constructed from FBLL×Hi-II (Lowe *et al*., 2006). Armstrong et al. used a low frequency of EC initiation line B73 to backcross a high tissue culture response line A188, generating the BC_6_S_4_ lines with high frequency of EC initiation (Armstrong *et al*., 1992). Five introgressed A188 segments were identified correlating with EC formation (Armstrong *et al*., 1992), and 4 markers located in or near the introgressed A188 segments were found involving EC formation using an F_2_ population of A188×Mo17 (Armstrong *et al*., 1992). Similarly, a B73 Near isogenic line WCIC2 (Donor parent: A188) with high frequency of EC initiation was used to genetically fine-map the QTLs for EC response, finally, a QTL located within a 3.06 Mb region on chromosome 3 was identified to control EC formation and regeneration (Salvo *et al*., 2018). Through reverse genetics, two genes, *ZmWUS2* and *ZmBBM*, were proved to regulate maize EC formation and regeneration. Overexpression of the two genes resulted in the improved frequencies of EC induction and transformation in both immature embryos and mature explants of the inbred lines with low tissue culture response (Lowe *et al*., 2016). However, few of genes responsible for tissue culture response were cloned in maize.

Maize shows remarkable genomic diversity among various inbred lines (Buckler *et al*., 2006; Lai *et al*., 2010; Schnable *et al*., 2009; Springer *et al*., 2009). The temperate line A188 (Gacheri *et al*., 2015), with high frequency of EC initiation (Armstrong *et al*., 1992; Salvo *et al*., 2018), is a desirable material to study the molecular mechanism underlying EC formation and regeneration. Due to the difference between A188 genome and B73 reference genome, these identified QTLs for embryo culture response have not been cloned so far, limiting their applications in improving EC formation capability. In addition, A188 shows considerable phenotypic variations from other inbred lines, such as plant height (Peiffer *et al*., 2014), ear height, days to tassel (Peiffer *et al*., 2014), days to silk (Peiffer *et al*., 2014), oil concentration (Cook *et al*., 2011), protein concentration (Cook *et al*., 2011), starch concentration (Cook *et al*., 2011) etc. (Table 1). Collectively, the assembly of a high-quality A188 reference genome is helpful to reveal the molecular mechanism underlying EC induction and other agronomic traits. Here we combine single-molecule sequencing and BioNano optical-mapping technologies to produce a *de novo* assembly of the A188 genome and provide these research communities with an excellent resource.

**Table 1.** The phenotypic performances of agronomic traits among different inbred lines

| Line | ECIR<sup>#</sup> | Sig. | Plant height* | Sig. | TBN* | Sig. | Ear No.* | Sig. | Pro con.<sup>#</sup> | Sig. | DtT | DtP | DtS |
| --- | --- | --- | --- | --- | --- | --- | --- | --- | --- | --- | --- | --- | --- |
| A188 | 91.53%±5.55% | a | 126.00±16.83 | d | 12.36±2.71 | a | 1.38±0.62 | a | 11.98±0.04 | b | 51 | 59 | 61 |
| W22 | 1.67%±1.85 | c | 159.50±17.03 | c | 11.48±2.48 | a | 1.29±0.60 | ab | 10.48±0.07 | c | 67 | 70 | 70 |
| Mo17 | 6.94%±1.39 | b | 184.29±17.99 | b | 6.21±1.05 | b | 1.07±0.68 | bc | 12.59±0.11 | a | 70 | 71 | 72 |
| B73 | 0±0 | c | 203.36±13.92 | a | 6.64±1.23 | b | 0.92±0.51 | c | 9.34±0.05 | d | 71 | 72 | 73 |

The results are shown as the means ± SD (“\*” n = 42, “#” n=3); “a, b, c, d” represents the significant
differences of trait with $P < 0.05$ . ECIR: embryonic callus induction ratio; Sig.: significance; TBN:
Tassel branch numbers; Pro con.: protein concentration; DtT: Days to Tassel; DtP: Days to Pollination;
DtS: Days to Silk.

## Methods

### Phenotypic evaluations of maize inbred line A188, B73, Mo17 and W22

The maize (*Zea Mays* L.) inbred lines A188, B73, Mo17 and W22, provided by Maize Research Institute of Sichuan Agricultural University, were grown in Chengdu (Sichuan province, China, N30°67’, E104°06’) in 2018. All of the lines were planted in a randomized complete block design with three replicates and two rows per line. A total of 14 plants were contained per row with the row length of 3 m and the row ledge of 0.75 m. These materials were managed according to the standard cultivation practices. At 10 days after pollination (DAP), the plant height and TBN were measured as described previously (Brown *et al*., 2011). The day duration from seeding to half of the plants tasseled, pollinated and silked was recorded as days to tassel, days to pollination and days to silk, respectively. The ear numbers were counted at 30 DAPs. Three mature seeds of each inbred line were crushed and subjected to the measurement of the protein concentration using the RAPID N exceed® (Elementar, Straße 1 63505 Langenselbold, Germany) according to the manufacturer’s instructions.

Maize inbred lines A188, B73, Mo17 and W22 were planted in a greenhouse (14/10 h light/dark, at 28°C and 70% relative humidity). Twelve days after self-pollination, 108 immature embryos (with 1.2-1.5 mm in length) from each line were collected, and evenly distributed among three Petri dishes containing the modified N6 medium (Frame *et al*., 2002) with scutellum upward to induce EC, with three replicates. After aseptic incubation for 21 days in darkness at 28°C, we investigated the EC induction ratio which was represented by (number of the immature embryos successfully inducing EC/ number of inoculated immature embryos) × 100%.

### Genomic DNA and total RNA isolation

The plants of A188 were grown in a greenhouse at 28 □ in a dark condition for 14 d. The yellow leaves of A188 seedlings were isolated and frozen immediately in liquid nitrogen for extracting genomic DNA, which was subsequently used for constructing libraries for PacBio sequencing and BioNano optical maps. To assist gene annotation and transcriptome analysis, transcriptome of five tissues were performed single-molecule long-read sequencing.

Total RNA was extracted from five tissues (12-d seedlings, tassel, silk, pericarp and 20-DAP seeds) with TRIzol reagent (Invitrogen, Carlsbad, CA) according to the manufacturer’s instructions. For each sample, we generated three independent biological replicates. The same amount of RNAs (1 μg) for each replicate of each tissue were pooled and stored at -80 □.

### PacBio library construction and sequencing

Libraries for SMRT PacBio genome sequencing were constructed as previously reported (Pendleton *et al*., 2015). Briefly, 20 μg of high-quality genomic DNA was sheared, and the ∼20 kb targeted size fragments were selected for ligation with SMRT adapter, followed by purification and size selection with Agilent 2100. The obtained PCR-free SMRTbell libraries were sequenced on the PacBio Sequel platform (Pacific Biosciences).

One microgram of enriched polyA RNA was reversely transcribed into cDNA by using the Clontech SMARTER cDNA synthesis kit, which was subjected to size selection using the BluePippin system. Size fractions eluted from the run were re-amplified to generate 2 libraries (0-1 kb and 1-10 kb). Then 2 μg cDNA of each library were subjected to Iso-Seq SMRTBell library construction according to the protocol reported on the website (https://pacbio.secure.force.com/SamplePrep). The SMRTBell libraries were then subjected to single-molecule sequencing on the PacBio Sequel platform (Pacific Biosciences).

### Optical library construction and sequencing

The professional kit was used to extract the High Molecular Weight (HMW) genome by agarose-embedded cells followed by protein digestion. The HMW genome of the quality check-through was specifically recognized by the *BspQ* I enzyme to identify the site to be labeled, and the fluorescent group is added to the double-stranded DNA molecule by means of modification and labeling, thereby ensuring the stability of the double strand and increasing sequence tag density at the same time. Finally, DNA molecules with fluorescent tags are stained to complete genome-specific tagging.

The marked library was loaded on the Irys chip for scanning and photographing. During the whole process, images were continuously converted into map data. After the real-time statistics and quality of Access reached the standard, the instrument was stopped and the data was transferred to bioinformatics analysis.

### *De novo* assembly of PacBio SMRT reads

After removing the short polymerase reads, low quality polymerase reads and self-connected adaptor sequences, 27,248,178 subreads (approximately 224G) were used for contig assembly with Falcon (Pendleton *et al*., 2015). Firstly, all of the subreads were pairwise compared to correct the error sequence with the parameters ‘--length_cutoff 12000 --length_cutoff_pr 14000’, followed by preliminary assembly with parameters: --min_idt 0.70 --min_cov 2 --max_n_read 200, and further error correction. The overlap graphs were constructed with parameters ‘--max_diff 100 --max_cov 100 --min_cov 2’, and then the contigs were assembled based on it. The Blasr (Chaisson and Tesler, 2012) was employed to map all the subreads back to the contigs with the parameters ‘--bestn 1 --maxScore -1000 --hitPolicy randombest’. We further performed assembly error corrections using Arrow (https://github.com/PacificBiosciences/GenomicConsensus/) with default parameters.

### Construction of BioNano optical maps

High-molecular-weight DNA was digested by the endonuclease BspQI and then labeled with IrysPrep Labeling mix and Taq polymerase according to standard BioNano protocols. The BioNano raw data was filtered using Bionano Access (version 1.0.3), generating high quality data. Then the BioNano Irys system was subsequently used to automatically image the labeled DNA. IrysSolve (https://bionanogenomics.com/support/software-downloads/) was used to *de novo* assemble the BioNano bnx files into genome maps. The RefAligner (https://bionanogenomics.com/support/software-downloads/) was used for molecule Pairwise comparison to identify overlaps, followed by construction of consensus maps. We recursively refined and extended the consensus maps by mapping all molecules back to the consensus maps.

### Hybrid assembly of PacBio contigs and BioNano optical maps

The PacBio-assembled contigs and BioNano-assembled genome maps were subjected to hybrid assembly by using the ‘HybridScaffold’ module of the IrysSolve as described previously (Sun *et al*., 2018). Briefly, the PacBio genome maps were aligned to an in silico BspQI-digested cmap. The BioNano genome maps were then aligned to the PacBio genome maps with RefAligner, followed by identifying and resolving the conflict points. After resolving the conflict points, Bionano genome maps and PacBio genome maps were merged to generate hybrid scaffold. The PacBio genome maps were mapped to hybrid scaffolds again to identify overlaps. If the overlap between PacBio contigs and hybrid scaffold was longer than 1 kb and identity ≥ 95%, these two sequences were merged. Based on the alignment information, the super-scaffolds were built.

### Construction of pseudomolecules

The A188 scaffolds were mapped to B73_RefGen_v4 genome using Bwa (version bwa-0.7.15, http://bio-bwa.sourceforge.net/bwa.shtml). The mapping rate of each scaffold mapped to each chromosome of B73 were calculated. The scaffolds with the highest mapping rate were kept to determine mapping to which B73 chromosome. The mapping file was further filtered according to the one-to-one correspondence, and the comparison noise was excluded. The remaining mapping results were submitted to combining coordinate to determine the alignment of scaffold on the B73 chromosome. Reverse alignments of the scaffolds were also performed based on the filtered mapping result file, and finally the scaffolds were anchored to the B73 reference genome according to the filtered position alignment information.

### Assembly evaluation

BUSCO (Benchmarking Universal Single-Copy Orthologs: http://busco.ezlab.org/) combined with tblastn, augustus, and hmmer softwares were used to evaluate the genome-assembly completeness. ‘Embryophyta_odb9’ that contained 1,440 single-copy orthologous genes was used as a searching dataset and was employed to assess the assembly completeness of the A188 genome.

### Repetitive elements prediction

TRF v4.07b (http://tandem.bu.edu/trf/trf407b.linux64.download.html) was used to predict tandem repeat. LTR Finder (Xu *et al*., 2010), RepeatScout (v1.0.5, http://www.repeatmasker.org), and PILER (v1.0, http://www.drive5.com/piler) were used to predict LTR element, LINE, SINE, and transposable DNA, respectively. First of all, the low complexity and low copy results of RepeatScout and PILER were removed. The predicted repetitive element sequences longer than 100 bp and the gap length ≤ 5% were kept and further mapped to protein sequences in SwissProt (ftp://ftp.ebi.ac.uk/pub/databases/uniprot/knowledgebase/uniprot_sprot.fasta.gz), the sequence alignments with non-transposable element protein sequences with evalue ≤ 1e^-4^, identity ≥ 30, coverage ≥ 30%, and length ≥ 90 bp were removed. Then the remained sequences were aligned to Rfam 11.0 database (ftp://ftp.ebi.ac.uk/pub/databases/Rfam) using BLASTN to remove ncRNA, and the predicted repetitive elements with evalue ≤ 1e^-10^, identity ≥ 80, and coverage ≥ 50% were removed. Moreover, the remained repetitive elements were aligned to RepBase and TE protein database using WU-BLAST, and were classified using RepeatClassifier, with the known simple repeat, satellite, and ncRNA sequences removed. The remained repetitive elements were compared to each other using BLASTN, the sequences with evalue ≤ 1e^-10^, identity ≥ 80, coverage ≥ 80%, and mapping length ≥ 80 bp were removed. Finally, the interspersed repeats were generated by masking predicted repetitive elements, known repetitive elements (RepBase), and protein repeat sequence (TE protein database) using RepeatMasker, RepeatMasker, and RepeatProteinMask, respectively.

### Gene annotation

MAKER2 (http://www.yandell-lab.org/software/maker.html) (Cantarel *et al*., 2008) was used to annotate genes in the A188 genome with the strategy as described previously (Sun *et al*., 2018). First, for protein-homology-based prediction, we downloaded the proteins of B73 reference genome, Mo17 reference genome, and W22 reference genome from gramene (http://gramene.org/) (Tello-Ruiz *et al*., 2015) as input of MAKER2. The A188 transcripts assembled from five different tissues based on single-molecule long-read sequencing in this study, B73 full-length transcripts from Iso-seq (Wang *et al*., 2016), and Mo17 transcripts (Sun *et al*., 2018) were used for gene transcript prediction. Second, the generated gene models were submitted to Augustus (Keller *et al*., 2011), SNAP (http://snap.stanford.edu/snap/download.html), GeneMark-ESSuite (version 4.32 http://topaz.gatech.edu/GeneMark/license_download.cgi), and Glimmerhmm (http://ccb.jhu.edu/software/glimmerhmm/) ab initio prediction softwares to further *de novo* predict gene models. Then, we further filtered the preliminary prediction gene set according to AED scores generated in MAKER software and the high confidence gene models generated finally.

### Identification of PAV sequences

To identify presence/absence-variation sequences (PAV, length longer than 500bp), we used a sliding-window method as reported previously (Sun *et al*., 2018). To identify A188-specific PAV sequences to B73, the A188 genome was divided into 500 bp windows with a step size of 100 bp. Then all of the 500 bp windows were aligned to B73 genome with BWA mem (Li, 2013) (http://bio-bwa.sourceforge.net/) with options ‘-w 500 -M’. The A188-sepecific PAV sequences are the sequences that cannot be aligned to the B73 genome or the primary alignment coverage less than 25% (Sun *et al*., 2018). Two overlapped PAV windows were merged. The same method was used to identify A188-specific PAV sequences to Mo17 and to W22, B73-specific PAV sequences to A188, Mo17-specific PAV sequences to A188, and W22-specific PAV sequences to A188. The PAV sequences within 100 kb of the physical coordinates were further merged. The merged region had more than 10% PAV sequences were defined as a PAV cluster. Finally, all of the PAVs were anchored back to corresponding genome. We used the same method (Sun *et al*., 2018) to identify A188-specific genes to B73, Mo17 and W22, respectively. In brief, the genes with more than 75% of the CDS regions falling in PAV sequences were defined as PAV genes.

### Comparative genomic analysis among A188, B73, Mo17 and W22

The Mummer software (Kurtz *et al*., 2004) was used to perform comparative genomic analysis between A188 and B73 genomes. Each A188 pseudo-chromosome sequence was mapped to the corresponding B73 chromosome using mummer with the parameters “nucmer -g 1000 -c 90 -l 40”. The mapping results were submitted to delta-filter to perform noise filtration with parameters “-r -q”. The show-coords was used for conversion of aligned physical coordinates with parameters “-rcloTH”. SNPs and small InDels (<100 bp) were identified using show-snps with “-ClrTH”. Show-diff was employed to obtain Inversions with the default parameters. Finally, we further filtered the alignments with aligned physical positions in one genome that were located more than 10 Mb away in another genome. The comparative genomic analysis between A188 and Mo17, A188 and W22 genomes were processed with the same method.

## Results

### Remarkable phenotypic difference between A188 and the other assembled lines

As a public inbred line, A188 has an outstanding response to the tissue culture, generating an approximate 100% efficiency in forming EC from immature embryos (Hodges *et al*., 1986; Ishida *et al*., 1996). However, previous studies demonstrated that B73 and Mo17 both have a very low frequency of inducing EC under standard conditions (Frame *et al*., 2006; Salvo *et al*., 2018). We also compared the phenotype of EC formation ratio among A188, B73, Mo17 and W22, indicating the highest EC induction ratio of A188 and the low EC induction ratios of the other three lines (Table 1). Moreover, the other agronomic traits show significant difference between A188 and the other lines including plant height, tassel branch numbers, ear numbers, protein concentration, days to tassel, days to pollination, and days to silk (Table 1 and Fig. 1). Our findings were in agreement with the previous studies about the phenotypic performances of A188 (Cook *et al*., 2011; Peiffer *et al*., 2014), implicating that A188/B73, A188/Mo17 and A188/W22 are therefore the ideal pairs of maize lines in genetic and molecular studies of these traits.

**Fig. 1.**
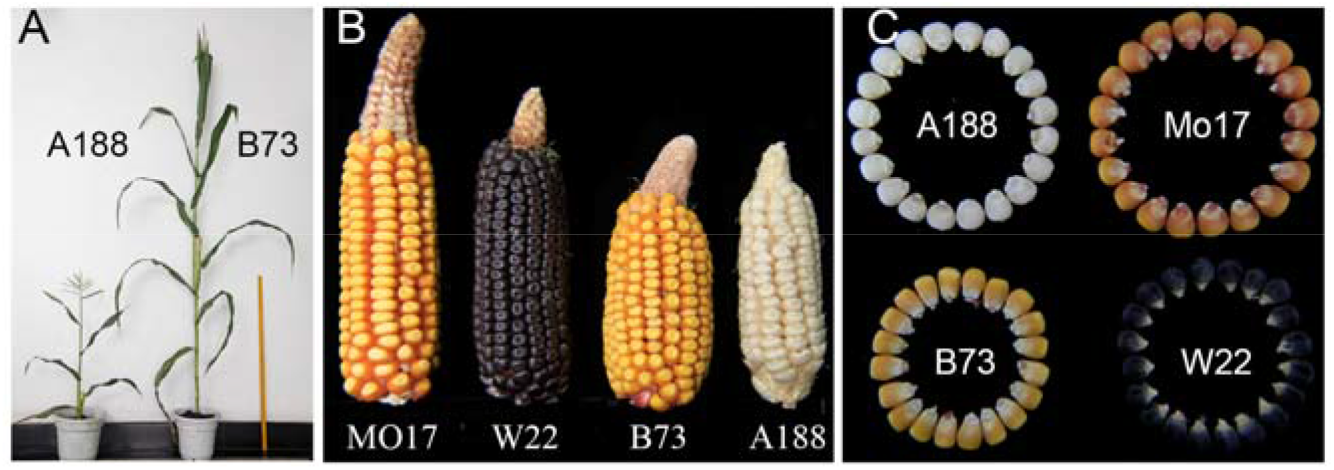
Overview of the trait difference between A188, B73, Mo17 and W22 inbred lines, including plant height (**A**), ear trait (**B**), and kernel size (**C**).

### Genome sequencing and *de novo* assembly

By combining with optical genome mapping with the BioNano Genomics Irys System, PacBio Sequel platform was used to sequence and *de novo* assemble of the A188 genome. Firstly, over 104-fold coverage of sequence data (224.03 Gb in total) generated from PacBio Sequel technology was used to perform initial assembly, resulting in a 2127.72 Mb assembly with a contig N50 size of 1.06 Mb and the longest contig of 4.97 Mb (Table 2, Tables S1 and S2). We then used 631.48-Gb BioNano molecule (287-fold-coverage BioNano optical map) to scaffold the assembled contigs and generated the final assembly which contains 4,469 scaffolds with a scaffold N50 size of 11.61 Mb and the longest length of 47.84 Mb (Table 2 and Tables S1 and S2). The total size of the final assembly was 2,207.74 Mb, similar to those of the B73 (2,106 Mb) (Jiao *et al*., 2017), Mo17 (2,183 Mb) (Sun *et al*., 2018) and SK (2,094 Mb) (Yang *et al*., 2019) genomes (Table 2 and Table S2). Bwa-0.7.15 was then used to anchor the A188 scaffolds to ten pseudo-chromosomes based on the B73 RefGen v4 genome according to the filtered position alignment information (Methods). Finally, 295 scaffolds were anchored and oriented onto ten chromosomes (2,084.35 Mb, 94.30% of the final genome assembly) and 3704 scaffolds failed to be mapped to chromosomes (5.70% of the final genome assembly) (Table S3). The final A188 assembly had 2,480 gaps (89.56 Mb in length), compared with 2,522 gaps in B73 and 238 gaps in SK genome (Yang *et al*., 2019). BUSCO (Simão *et al*., 2015) was used to evaluate the A188 genome assembly quality. Finally, 95.3% of complete single-copy BUSCOs could be aligned to the A188 final assembly, similar to those for the B73 (Jiao *et al*., 2017), Mo17 (Sun *et al*., 2018), W22 (Springer *et al*., 2018) and SK (Yang *et al*., 2019) genomes, indicating the near completeness of our assembly (Table S4).

**Table 2.** Global statistics for the A188 genome assembly

|  | PacBio assembly | Pacbio+Bionano hybrid<br>assembly | Pseudomolecule |
| --- | --- | --- | --- |
| Total length of assembly (Mb) | 2,127.72 | 2,210.33 | 2,084.35 |
| N50 size (Mb) | 1.06 | 11.61 | - |
| Longest length (Mb) | 4.97 | 47.84 | - |
| Number of sequences | 6,385 | 4,469 | 10 |

### Genome annotation

A modified approach based on the annotation pipeline (Sun *et al*., 2018) was employed to analyze the transposable-element content of our A188 assembly. Finally, approximately 80.70% of the A188 genome sequence were identified as transposable-element sequences, including retrotransposons (71.93%), DNA transposons (5.91%), and unclassified elements (2.49%) (Table S5), which was lower than those in B73 (Jiao *et al*., 2017), Mo17 (Sun *et al*., 2018), W22 (Springer *et al*., 2018), SK (Yang *et al*., 2019) and K0326Y (Li *et al*., 2020) genomes. For retrotransposons, the families of *Copia* and *Gypsy* represented 24.01% and 46.92% of the A188 genome, respectively (Table S5). For DNA transposons, the family of *hAT* was much lower than those in the B73 (Jiao *et al*., 2017) and Mo17 (Sun *et al*., 2018) genomes.

To annotate the A188 genes, we integrated two technologies, protein-homology-based prediction and isoform sequencing of five different A188 tissues, and combined the reported B73 full-length transcripts (Wang *et al*., 2016) and Mo17 transcripts (Sun *et al*., 2018). In total, 44,653 high-confidence protein-coding gene models with 66,359 transcripts were predicted (Table S6). Among them, 10,965 (24.56% of the predicted genes) and 16,243 (36.38% of the predicted genes) genes were supported by ISO-seq with CDS coverage >90% and >50%, respectively (Table S6). In total, 41,715 (93.42%) of the predicted A188 genes were mapped to ten pseudo-chromosomes (Table S3). In addition, 93.52% (62,058) of these transcripts were functionally annotated and deposited in the public databases (Fig. S1).

### Genome structural variations between A188, B73, Mo17 and W22

To better understand the genome difference, we individually aligned the pseudo-chromosomes of B73, Mo17 and W22 to those of A188. In total, 62.50% (1,316.38 Mb), 63.10% (1,327.82 Mb) and 62.59% (1,327.48 Mb) of the B73, Mo17 and W22 genome sequences matched in one-to-one syntenic blocks with 63.16% (1,316.45 Mb), 63.71% (1,328.03 Mb), and 63.69% (1,327.43 Mb) of the A188 genome sequence, respectively (Fig. 2, Fig. S2, and Table S7).

**Fig. 2.**
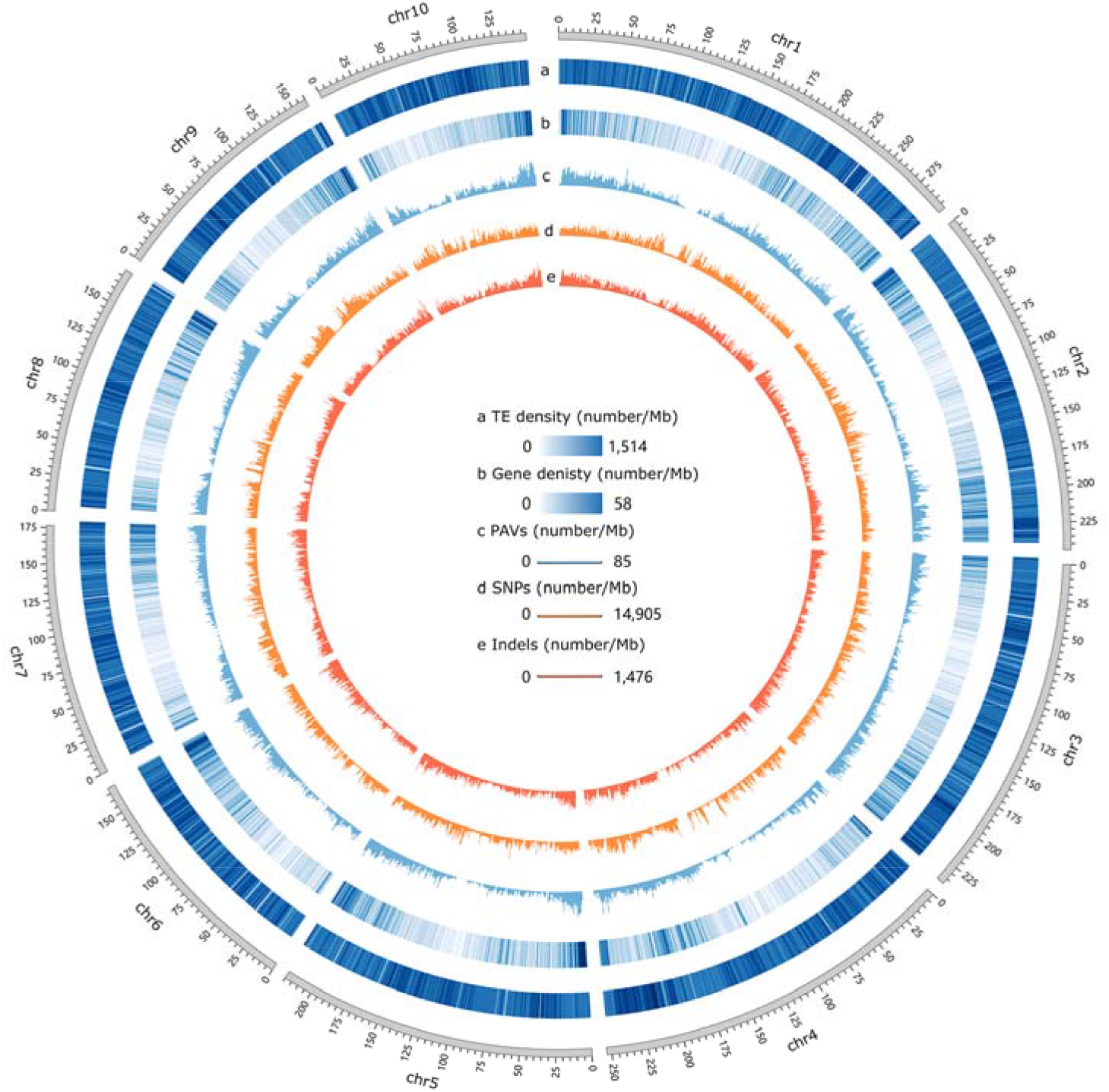
Features of the A188 genome. **a**, Transposable-element density; **b**, gene density; **c, d** and **e**, numbers of PAVs (c), SNPs (d) and InDels (e) compared with B73 genome. The sliding window is 1 Mb for all tracks.

A total of 9,865,320 SNPs, 634,693 insertions and 654,322 deletions were identified between A188 and B73, with an average of 4.73 SNPs, 0.30 insertions and 0.31 deletions per kilobase (Fig. 2, Fig. S3, and Table S7). We also identified 9,490,058 SNPs, 743,829 insertions and 654,841 deletions between A188 and Mo17, and 9,614,783 SNPs, 600,755 insertions and 628,504 deletions between A188 and W22 (Fig. S3, Table S7). Less than 2.5% of these variations in A188 are allocated in CDS regions, and the remainders are annotated as intergenic variations (Table 3 and Table S8). Interestingly, a genome-wide comparison showed that InDels of 3N±1 bp in CDS region were more abundant than 3N bp in gene coding regions (Table 3), between A188 and any one of lines B73, Mo17 and W22. We then focused on identifying presence/absence-variation sequences (PAV) longer than 500 bp in the A188 genome. By comparing the A188 and B73 genomes, we identified 27,240 A188-specific genomic segments (16.92 Mb in total) and 28,558 B73-specific genomic segments (17.76 Mb in total). Most of these PAV segments were shorter than 3 kb, only 1 and 2 PAV segments were identified longer than 3 kb in A188 and B73, respectively (Fig. S4). Similarly, by comparing the A188 and Mo17 genomes, we identified 26,983 A188-specific genomic segments (16.76 Mb in total), and 28,030 Mo17-specific genomic segments (17.44 Mb in total). Most of the PAV segments were shorter than 3 kb, only 1 and 3 PAV segments were identified longer than 3 kb in A188 and Mo17, respectively (Fig. S4). The comparison of the A188 and W22 genomes identified 31,536 A188-specific genomic segments (19.42 Mb in total), and 29,192 W22-specific genomic segments (17.98 Mb in total), with 1 and 4 PAV segments longer than 3 kb in A188 and W22, respectively (Fig. S4). According to the criterion that a gene with ≥ 75% of coding sequences covered by a PAV sequence can be identified as a PAV gene (Sun *et al*., 2018), we identified 100 A188-specific and 104 B73-specific PAV genes, by comparison of A188 and B73 genomes. Similarly, 116 A188-specific and 146 Mo17-specific PAV genes were found by comparing A188 and Mo17, and 112 A188-specific and 116 W22-specific PAV genes were identified between A188 and W22 (Table 3 and Table S9). Combined these findings illustrated that the A188 genome showed huge variations from B73, Mo17 and W22 genomes. However, only 9 A188-specific PAV genes were simultaneously identified in comparison with the other three inbred lines, thus illustrating that most of the A188-specific PAV genes have already existed in other lines (Table S9).

**Table 3.** Variations within genes between A188, B73, Mo17 and W22 genomes

| Variation type | A188 to B73 | A188 to Mo17 | A188 to W22 |
| --- | --- | --- | --- |
| 1. Structurally conserved genes | 30635 | 31095 | 30764 |
| No DNA variation in CDS | 20557 | 21007 | 20713 |
| No DNA variation in CDS and intron | 17168 | 17634 | 17382 |
| No DNA variation in genic region* | 8647 | 9054 | 8854 |
| Without amino acid substitutions | 23989 | 24424 | 20860 |
| With amino acid changes | 22958 | 23313 | 23070 |
| With missense mutation in CDS | 21975 | 21601 | 21869 |
| With 3N InDel in CDS | 7210 | 7257 | 7295 |
| 2. Genes with large structural mutations | 13020 | 13044 | 12939 |
| Start codon mutation | 737 | 747 | 742 |
| Stop codon mutation | 506 | 504 | 506 |
| Splice donor mutation | 841 | 801 | 811 |
| Splice acceptor mutation | 1671 | 1559 | 1486 |
| With 3N±1 InDel in CDS | 10834 | 10982 | 10889 |
| Premature termination codon | 2362 | 2355 | 2389 |
| 3. PAV genes | 204 | 262 | 228 |
| A188 present PAV genes | 100 | 116 | 112 |
| A188 absent PAV genes | 104 | 146 | 116 |
| Total of genes with large structural variations | 13224 | 13306 | 13167 |

\* Genic regions include the 2 kb upstream and downstream of the gene body.

### Gene structural variations

A total of 20,557, 21,007 and 20,713 genes displayed no variations in the CDS regions between B73 and A188, Mo17 and A188, and W22 and A188, respectively (Table 3). Moreover, 17,168, 17,634 and 17,382 A188 genes showed no variations in CDS and intron regions as compared with B73, Mo17 and W22, respectively (Table 3). In particular, as compared with B73, Mo17 and W22, 8,647, 9,054 and 8,854 genes were highly conserved without any genetic variation (including 2 kb upstream and downstream), respectively (Table 3). Moreover, we found 23,989, 24,424 and 20,860 A188 genes showed synonymous variations in CDS compared to B73, Mo17, and W22, respectively (Table 3). Compared with B73, 22,958, 21,975 and 7,210 genes in A188 resulted in amino acid changes, missense mutation and non-frameshift InDels, respectively (Table 3). Mapped to Mo17, 23,313, 21,601 and 7,257 genes in A188 were identified to contain amino acid changes, missense mutation in CDS and non-frameshift InDels, respectively (Table 3). Aligned to W22, 23,070, 21,869 and 7,295 A188 genes showed amino acid changes, missense mutation in CDS and non-frameshift InDels, respectively (Table 3). All of these genes were classified as structurally conserved genes between A188 and the other lines, which accounted for ≥ 68.61% of the annotated A188 genes and may function in basic physiological effects.

By comparing B73 and A188 genomes, we identified 737, 506, 841, 1,671, 10,834 and 2,362 genes in A188 that generated start codon mutations, stop codon mutations, splice donor mutations, splice acceptor mutations, frameshift InDel in CDS and premature termination codon mutations, respectively (Table 3). Similarly, 747, 504, 801, 1,559, 10,982 and 2,355 genes in A188 led to start codon mutations, stop codon mutations, splice donor mutations, splice acceptor mutations, frameshift InDels in CDS and premature termination codon mutations, respectively, as compared with Mo17 (Table 3). As well, 742, 506, 811, 1,486, 10,889 and 2,389 genes in A188 showed start codon mutations, stop codon mutations, splice donor mutations, splice acceptor mutations, frameshift InDels in CDS and premature termination codon mutations, as compared with W22, respectively (Table 3). In addition, 204, 262 and 228 PAV genes were identified between A188 and the lines B73, Mo17 and W22 genomes, respectively (Table 3). Finally, a total of 13,224 (29.62 %), 13,306 (29.80 %) and 13,167 (29.49 %) A188 genes had large structural variations (start- or stop-codon mutations, splice-donor or splice-acceptor mutations, frameshift mutations, premature termination codon mutations or PAV variations) as compared with B73, Mo17 and W22 genomes, respectively.

### A188 genome-based genetic dissection of tissue culture response

Recently, by using the F_3:4_ population derived from B73 and WCIC2 (a near isogenic line of B73 containing several A188 segments), a locus associated with embryogenic and regenerative capabilities of immature embryo was fine-mapped within a 3.06 M region (chr3:178772856-181826658) based on the B73 reference genome, suggesting that the genes harbored by the A188 segment caused the high callus formation ratio (Salvo *et al*., 2018).

To explore the candidate genes of embryonic callus induction, we aligned the 3.06 M B73 segment to the A188 genome and revealed a 3.89 M syntenic segment (Fig. 3) in A188. Within the 3.89 M segment, 51, 57, and 57 A188 genes were identified syntenic to B73, Mo17 and W22 syntenic segment, respectively (Table S10). Among them, 6, 14, and 6 genes showed large structural variation (LSV: premature termination codon, stop codon loss, frameshift deletion, or frameshift insertion) relative to B73, Mo17 and W22, respectively (Table S11), and 4 LSV genes in A188 were simultaneously identified in comparison with the other 3 lines (Table 4).

**Table 4.**
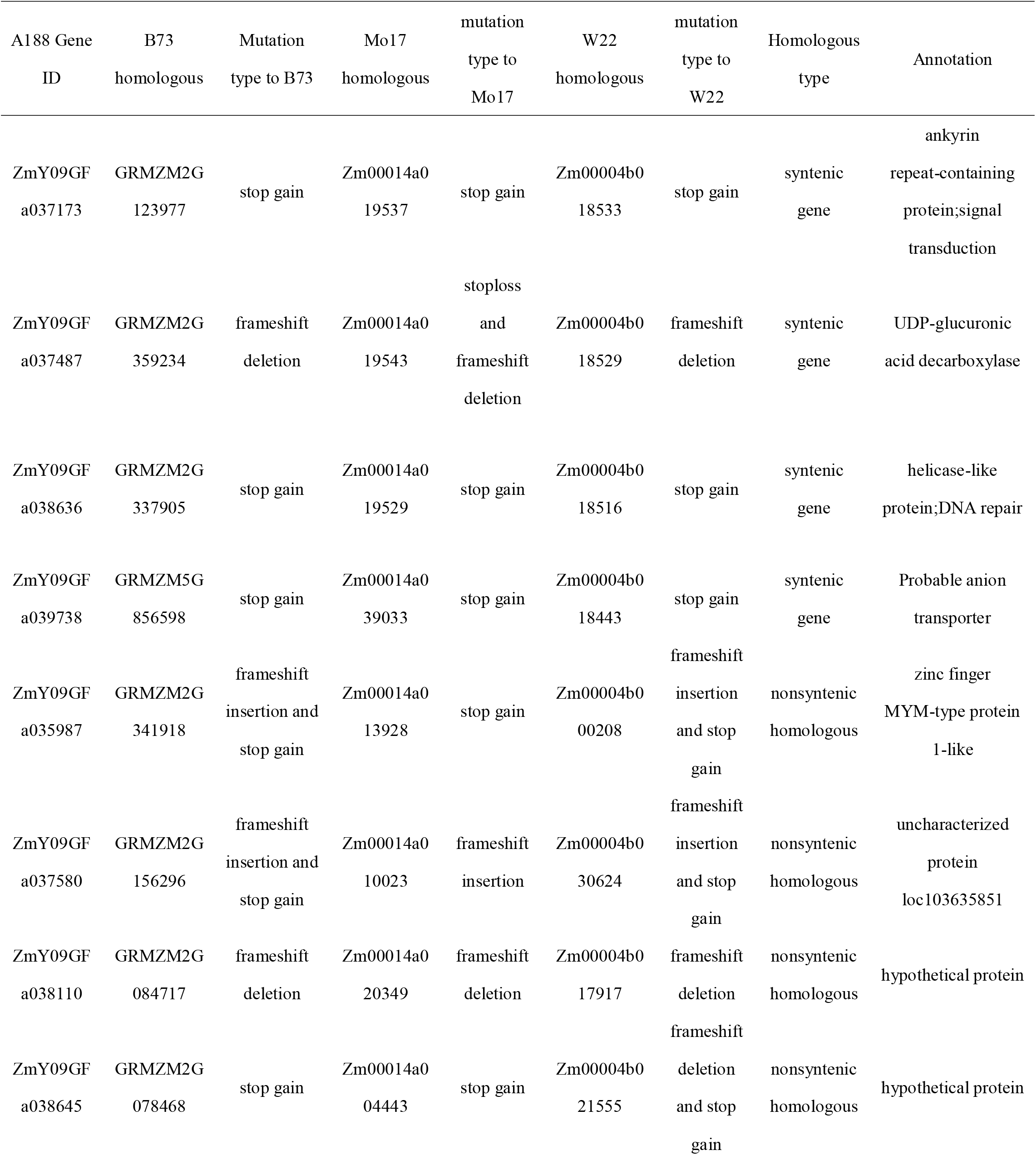
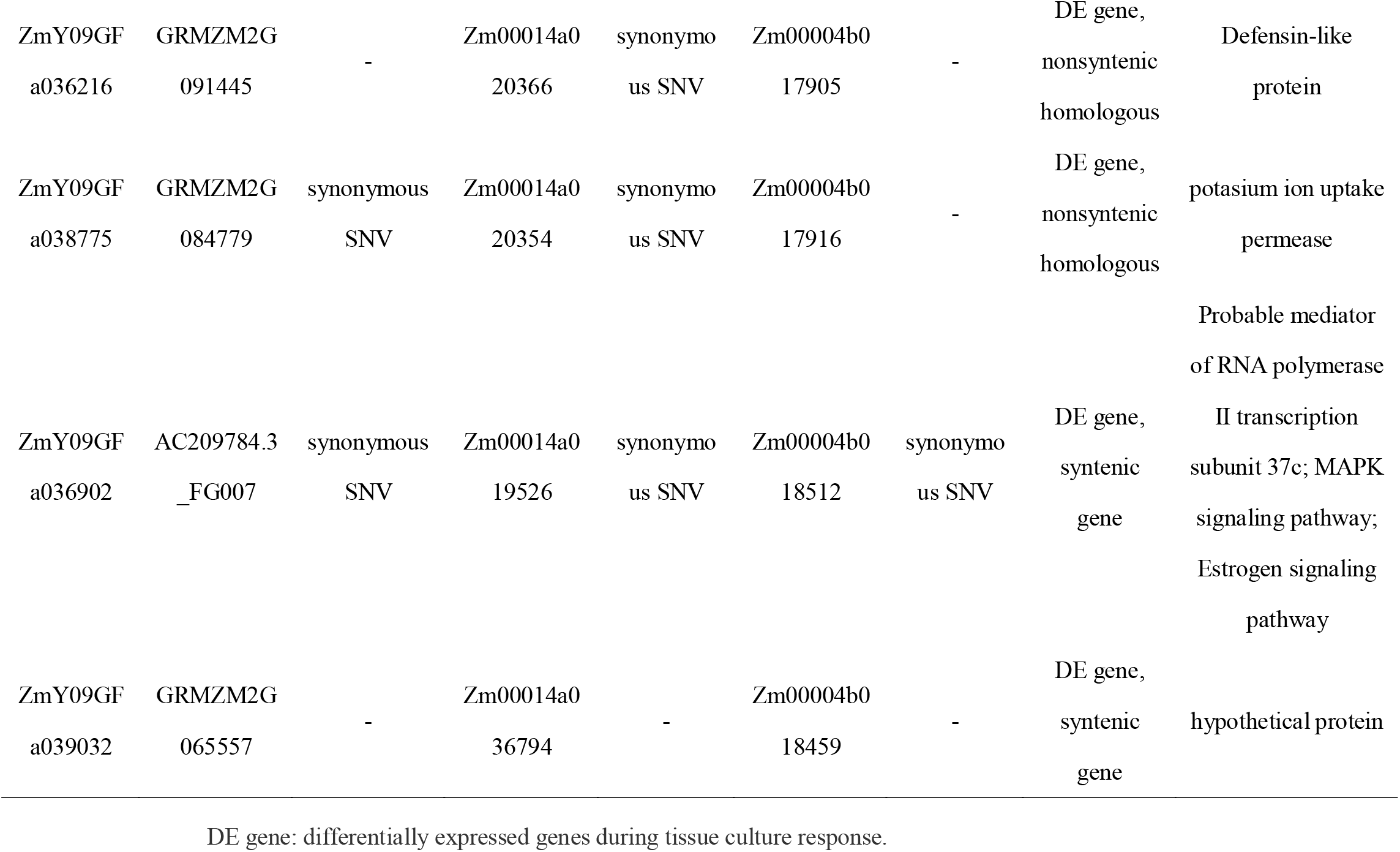
Tissue culture response candidate genes

**Fig. 3.**
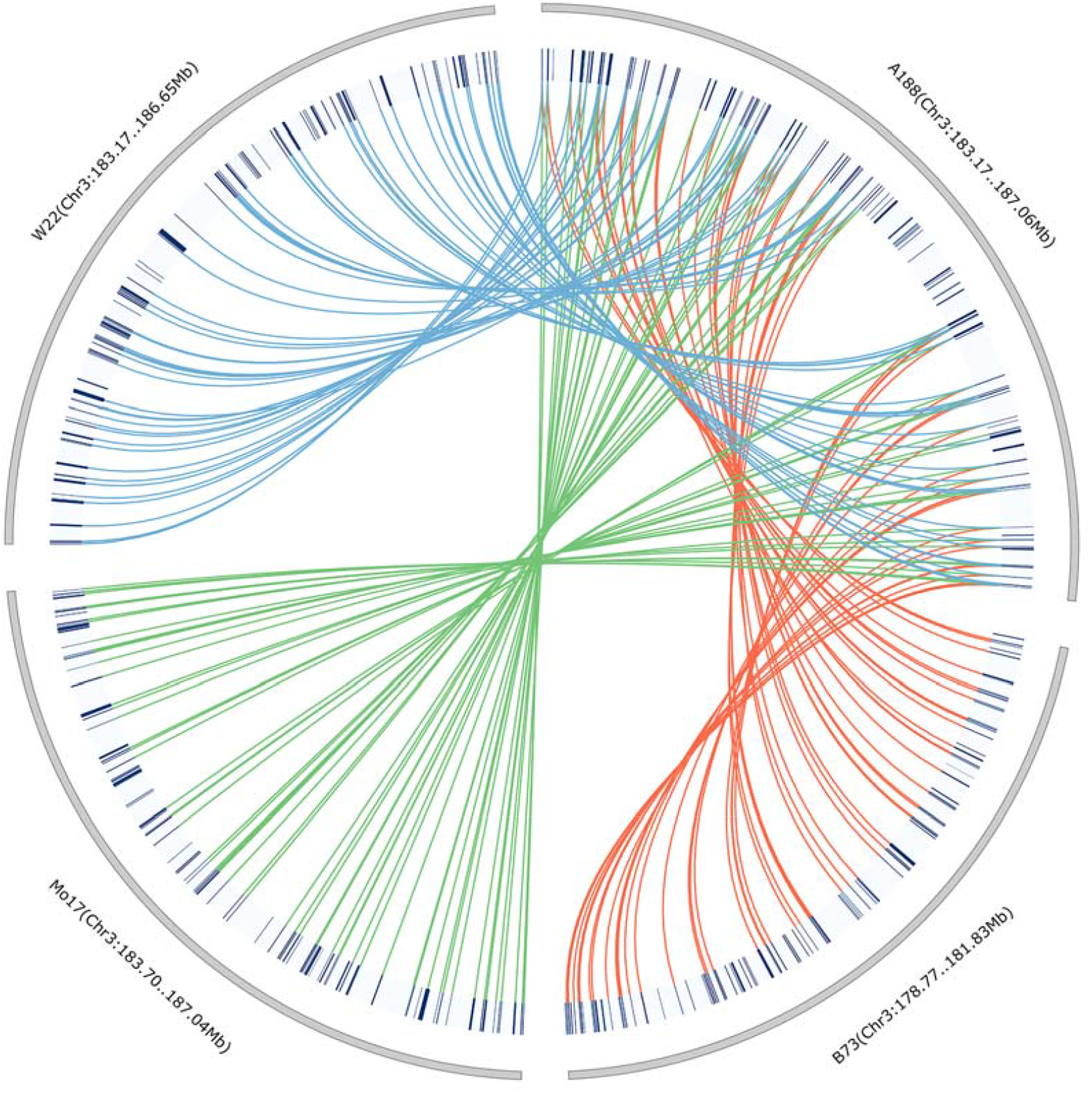
Tissue culture response candidate loci. The 3.89 M A188 segment (QTL for maize tissue culture response) aligned to the syntenic segment of B73, Mo17 and W22 genomes. The red, green and blue lines indicate aligned A188 genes in the 3.89 M segment to the B73, Mo17 and W22, respectively.

We also focused on the nonsyntenic genes of A188 in the QTL. Finally, we identified 48, 42, and 42 A188 genes in the QTL interval that were identified nonsyntenic, as compared with B73, Mo17 and W22 genomes, respectively (Table S10). To further identify whether the nonsyntenic genes have homologues in other sites of the 3 inbred lines, we mapped these nonsyntenic genes to the B73, MO17 and W22 whole genomes. Finally, 28, 11, and 24 A188 nonsyntenic genes showed LSV to their corresponding homologues in B73, Mo17 and W22, respectively (Table S11), and 4 LSV genes in A188 were simultaneously identified in comparison with the other 3 inbred lines (Table 4).

Moreover, previous studies have demonstrated that changes in gene expression can be induced during somatic embryogenesis to respond to tissue culture process (Ge *et al*., 2017; Salvo *et al*., 2018; Shen *et al*., 2012; Zhang *et al*., 2014). Based on the reported transcriptome data of A188 (Zhang *et al*., 2014), four of the 99 A188 genes within the mapped QTL were regulated by more than 8 folds in different stages of immature embryo culture, relative to control. Collectively, the 4 syntenic genes with LSV, the 4 nonsyntenic genes with LSV as well as the 4 differentially expressed genes were designated as the candidate genes responsible for tissue culture capability of A188 immature embryo in this study (Table 4).

## Discussion

Although A188 is limited from the application in breeding programs, due to its poor agronomic traits, A188 shows significant phenotypic variations from B73, Mo17 and W22, including plant height, tassel branch number, ear number, protein concentration, days to tassel, days to silk etc., especially EC induction ratio (Table 1). The phenotypic performance is determined by the combination of genotype and environment. To better understand the mechanisms underlying the phenotypic difference between A188, B73, Mo17 and W22, we sequenced and *de novo* assembled the A188 genome. Finally, we assembled the A188 genome into 2,207.74 Mb with a scaffold N50 size of 11.61 Mb (Table 2, Table S2). As expected, A188 showed large genomic variations as compared with B73, Mo17 and W22 (Fig. 2, Table 3, Tables S7, S8 and S9). Our new A188 genome provides a good resource to map causal genes controlling these various traits. We also identified a number of A188 genes presenting structure variations relative to other 3 inbred lines, such as genes with start/stop codon mutations, splice donor/acceptor mutations and frameshift InDels, which provides a novel view to study gene function and evolutionary analysis.

EC induction from maize immature embryo is highly dependent on genotype, which resulted in only a few functional genes identified. Accordingly, the molecular mechanism underlying EC induction still remains unclear in maize. Combined our new A188 genome, previously reported QTLs, and RNA sequencing data, we successfully identified 12 candidate genes responsible for maize tissue culture response (Table 4). These candidate genes provide new insight into understanding the molecular mechanisms of maize tissue culture response, and provide new gene resources for improving maize embryonic callus induction and maize genetic transformation, which will further contribute to gene function revelation and transgenic breeding in maize. Especially, ZmY09GFa037173 showed a premature termination mutation in A188, which was annotated as an Ankyrin repeat-containing protein and involved in signal transduction. In addition, the *Arabidopsis homologue, Itn1*, was previously reported to regulate ROS accumulation under salt-stress through regulating ABA signaling pathways (Sakamoto *et al*., 2008), which suggest that ZmY09GFa037173 have a potential to induce maize callus formation by mediating ROS levels.

Owing to the vast genetic diversity among maize germplasms, the currently identified genetic variants by comparison of nine public maize genomes are still unsaturated (Yang *et al*., 2019). The previously study suggest that more than 20 reference genomes of maize and teosinte were required for performing pan-genome construction, which will provide better coverage for genetic variations of the *Zea genus* (Yang *et al*., 2019). Our new sequenced and assembled A188 genome thereby provides a new reference genome and structure variation resource.

## Supplementary data

**Fig. S1**. Code gene function annotation result using the public databases of NR, Swiss-Prot, eggNOG, GO and KEGG.

**Fig. S2**. Whole-genome comparison of A188 versus B73 and Mo17. Grey lines represent the one-to-one aligned genes between each pair of pseudomolecules.

**Fig. S3**. Histogram of InDel number comparisons of A188 versus B73 and Mo17 of the whole genome and coding regions.

**Fig. S4**. Length distribution of PAV sequences between A188 and B73 genomes (a), A188 and Mo17 genomes (b), A188 and W22 genomes (c).

**Table S1**. Summary of sequencing data of A188 genome.

**Table S2**. Details of the A188 genome assembly.

**Table S3**. Details of the 10 A188 pseudo-chromosomes.

**Table S4**. BUSCO analysis.

**Table S5**. Comparisons of repetitive elements between A188, B73 and Mo17.

**Table S6**. Statistics of A188, B73 and Mo17 gene models.

**Table S7**. Summary of aligned sequences, SNPs and InDels in A188, B73, Mo17 and W22 genomes.

**Table S8**. Genome distribution of SNPs and InDels between A188, B73, Mo17 and W22 genomes.

**Table S9**. Summary of PAV genes in A188, B73, Mo17 and W22 genomes.

**Table S10**. Syntenic analysis of the 99 genes on the A188 syntenic segment compared with B73, Mo17 and W22 syntenic segments.

**Table S11**. Mutation type of the 99 genes within the QTL on the A188 syntenic segment compared with B73, Mo17 and W22 homologous genes.

## Acknowledgements

This study was supported by the National Natural Science Foundation of China (31871637 and 32072073), and the Project of Transgenic New Variety Cultivation (2016ZX08003003).

## Author contributions

Y.S., F.G. and L.M. designed the research. J.Q., P.L. and J.H. performed genome assembly, genome annotation and genome comparison. F.G. L.P. and J.H. prepared DNA/RNA samples and constructed the next-generation-sequencing library. P.L. performed phenotype investigation. Y.S., F.G., L.M., J.Q., P.L., L.P., J.H., C.Z., G.Y., C.Y., S.G., and G.P. participated in the analysis. F.G. J.Q., P.L., L.P., L.M. and Y.S. wrote and revised the manuscript.

## Data Availability

All datasets reported in this study have been deposited in GenBank (NCBI) with the following accession IDs: Genome assembly, JADZIA000000000; Raw data for genome assembly and gene annotation, PRJNA678284.

## Conflict of Interest Statement

The authors declare no conflict of interests.

## Notes

### Competing Interest Statement

The authors have declared no competing interest.

